# Evidence of Chitin in the Ampullae of Lorenzini of Chondrichthyan Fishes

**DOI:** 10.1101/687301

**Authors:** Molly Phillips, W. Joyce Tang, Matthew Robinson, Daniel Ocampo Daza, Khan Hassan, Valerie Leppert, Linda S. Hirst, Chris T. Amemiya

**Affiliations:** Department of Biology, University of Washington, Seattle, WA 98195, USA; Department of Molecular and Cell Biology, University of California, Merced, Merced, CA 95343, USA; Department of Orthopaedics and Sports Medicine, University of Washington School of Medicine, Seattle, WA 98109, USA; Department of Materials Science and Engineering, University of California, Merced, Merced, CA 95343, USA; Department of Organismal Biology, Uppsala University, 75236 Uppsala, Sweden; Department of Physics, University of California, Merced, Merced, CA 95343, USA

## Abstract

Chitin is synthesized by a variety of organisms using enzymes called chitin synthases and was recently discovered in a number of aquatic vertebrates. In our ongoing investigations into the presence of vertebrate chitin, we unexpectedly found evidence of the polysaccharide within the electrosensory organs, known as Ampullae of Lorenzini, of diverse chondrichthyan fishes. Experiments with histochemical reagents, chemical analyses, and enzymatic digestions suggested that chitin is a component of the hydrogel filling the structures. Further, *in situ* hybridization with a sequence from the little skate (*Leucoraja erinacea*) revealed that chitin synthase expression is localized to cells inside the organs. Collectively, these findings suggest that chondrichthyan fishes endogenously synthesize chitin and beg further investigation into the function of chitin in the electrosensory system.

## INTRODUCTION

We previously reported that the polysaccharide, chitin, a key component of arthropod exoskeletons and fungal cell walls, is endogenously produced by fishes and amphibians in spite of the widely held view that it was not synthesized by vertebrates [1]. Genes encoding chitin synthase enzymes were found in the genomes of a number of fishes and amphibians and shown to be correspondingly expressed at the sites where chitin was localized [1, 2]. In this report, we present evidence suggesting that chitin is prevalent within the specialized electrosensory organs of cartilaginous fishes (Chondrichthyes). These organs, the Ampullae of Lorenzini (AoL), are widely distributed but typically concentrated toward the rostral end of the fishes and comprise a series of gel-filled canals emanating from surface pores in the skin (Fig. 1A). The canals extend into bulbous structures (alveoli) containing sensory cells capable of detecting subtle changes in electric fields (Fig. 1B) [3, 4]. These findings extend the number of vertebrate taxa where endogenous chitin production has been detected and raise many questions regarding chitin’s potential function within chondrichthyan fishes and other aquatic vertebrates.

**Figure 1.**
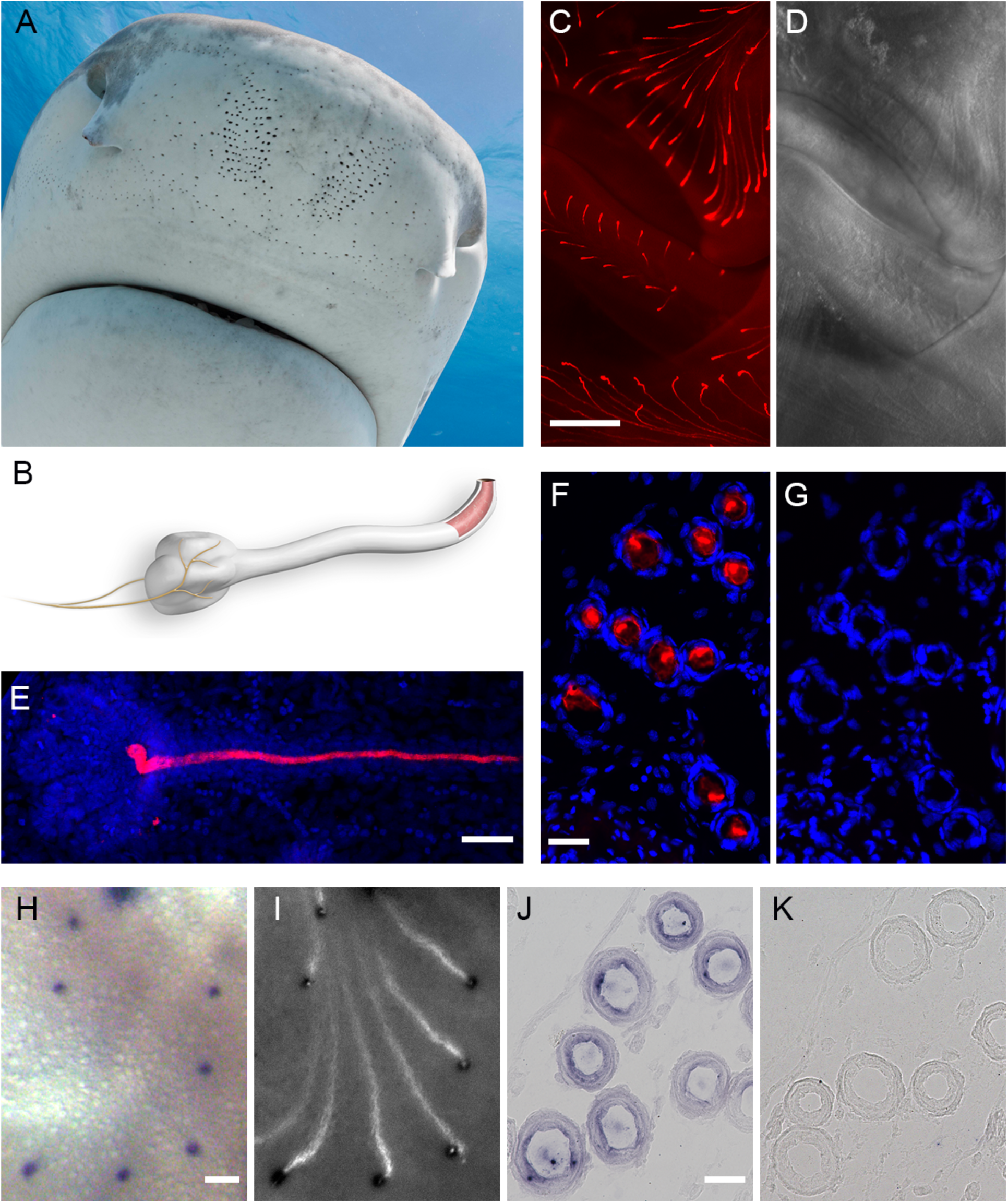
Description of Ampullae of Lorenzini and characterization of AoL chitin. (A) Snout of a tiger shark (*Galeocerdo cuvier*) revealing its numerous AoL pores. Photo taken by Neil Hammerschlag. (B) Illustration depicting an individual AoL. The pore at the upper right leads into a canal encased by a collagenous sheath filled with an acellular hydrogel (red) as depicted by the cutaway region. At the distal end of the organ is a round, lobed structure, the alveolus. The alveolus contains hundreds or thousands of electrosensory cells that synapse with afferent neurons and transmit information to the brain and which, collectively, make a topographical image of the surrounding electrical environment. (C, D) Whole-mount preparation of an embryonic little skate (*Leucoraja erinacea*) labeled with a fluorescent (red) chitin binding domain probe (C) and the corresponding brightfield image of the same specimen (D). The fluorescence pattern directly correlates with the distribution of AoL (raised canals on the surface of the skin). This field is the ventral surface around the lower lip (the middle band of AoL in C). (E) Confocal image of a chitin-labeled AoL from an embryonic catshark (*Scyliorhynus canicula*). Nuclei are stained with DAPI (blue) and delineates the canal and alveolus, and chitin (red) is seen throughout and across the canal. (F, G) *in situ* chitinase assay of AoL of embryonic *L. erinacea*. (F) Adjacent paraffin cross-sections of wing tissue were stained for chitin (red) after treatment with buffer (F) or chitinase (G), then counterstained with DAPI (blue). Chitin is seen within the AoL lumen of buffer-only controls (F) but not the chitinase-digested sample (G). (H-K) Confirmation of chitin synthase gene expression in the AoL by *in situ* hybridization (ISH). A skate antisense riboprobe (*LeCHS*) was used for ISH on whole-mount (H, I) or paraffin sections (J) of embryonic *L. erinacea*. The riboprobe (blue) localized to regions in the vicinity of the pores of the AoL (H), which were corroborated by CBD histochemistry (white) of the same tissue specimen (I). (J) Chitin synthase gene expression is seen in the endothelial cells lining the canal. (K) ISH negative control. Scale bars are 500 μm (C, D), 50 μm (E), 20 μm (F, G), 50 μm (H, I), and 10 μm (J, K).

## RESULTS

Affinity histochemistry was carried out with fluorescent probes containing a chitin binding domain (CBD) cloned from a bacterial chitinase enzyme [1]. Experiments using formalin-fixed little skate (*Leucoraja erinacea*) embryos revealed an overt correspondence of CBD signals with the AoL (Fig. 1C,D). In experiments using confocal microscopy, CBD signal was observed throughout the AoL (Fig. 1E), specifically localized to the hydrogel filling the acellular lumen of the organs (Figs. 1F, S1A). The space-filling nature of AoL gel within AoL canals of a juvenile shark and adult skate can be visualized in Fig. S1A and B, respectively.

These initial findings led us to investigate the potential existence of AoL chitin using multiple methods. First, we used chitin-digesting enzymes (chitinases) on AoL-containing histological sections. When chitinase was applied to the sections, we observed the elimination of detectable CBD signals (Fig. 1G) as compared to controls (Fig. 1F). This was done with AoL-containing tissues from multiple chondrichthyan species. Further, in addition to the bacterial chitinase CBD, we used another probe, containing an unrelated chitin binding domain, on whole-mount and histological preparations of a few chondrichthyan species, including spotted ratfish (*Hydrolagus colliei*) (Fig. S1D,E). Both probes yielded highly similar patterns despite their disparate amino acid sequences [1].

We used antisense riboprobes specific to a chitin synthase from *L. erinacea* (*LeCHS*) for *in situ* hybridization (ISH) on both whole-mount (Fig. 1H,I) and sectioned tissues (Fig. 1J,K). The whole-mount ISH patterns using the *LeCHS* riboprobe (Supplement) indicated expression at the AoL pores (Fig. 1H), and that CBD signal corresponded with these sites (Fig. 1I). The ISH signals in this particular experiment were restricted to the pores, however, we have observed that probe penetration can be variable likely due to the fish’s thick skin. We thus used sectioned materials and found *LeCHS* expression in the epithelial cells lining AoL canal walls at various canal depths (Figs. 1J, S1C). The specific co-localization of *LeCHS* expression and fluorescent CBD signals is compelling evidence that CBD is likely binding to chitin, not some other related polysaccharide, and that the epithelial cells within the AoL are producing and secreting chitin. Additionally, the ISH signals shown in Fig. S1C, which are cross-sections of AoL at different levels and transverse angles, indicate that cells inside the alveoli do not exhibit appreciable chitin synthase gene activity.

Lastly, to investigate whether or not the Ampullae of Lorenzini contain chitin *per se*, we chemically extracted polysaccharides from spotted ratfish (*Hydrolagus colliei*) gel for analysis by Fourier transform infrared spectroscopy (FTIR), atomic force microscopy (AFM), scanning electron microscopy (SEM), and monosaccharide analysis. The resultant FTIR spectrum contained peaks characteristic of two different allomers of chitin, sharing the most similarities with β-chitin from cuttlebone (Fig. S2A). Further, when the extracted polysaccharides were imaged with SEM and AFM, crystals reminiscent of chitin nanowhiskers were readily observed (Fig. S2B-E) [5]. A preponderance of glucosamine in the monosaccharide chemical analysis of this material is consistent with the structure of homopolymeric chitin (Fig. S2F,G).

Throughout our studies, we observed the localization of CBD to the AoL of diverse chondrichthyan species belonging to both elasmobranch and holocephalan lineages (Fig. S1F). This could suggest that the use of chitin in electrosensory organs evolved in the ancestor to all cartilaginous fishes, perhaps in the placoderms [6]. A more detailed treatment of chitin synthase genes within Chondrichthyes will be reported elsewhere.

The Ampullae of Lorenzini are sensory organs that detect changes in weak electric fields at the skin surface and transduce them into neuronal signals that project onto the brain. The subsequent topographical map of the organism’s electrical surroundings is thought to be used for various biological functions, most notably the detection of prey, localization of conspecifics for mating, and navigation [3]. While the electrophysiology of voltage-gated ion channels in the alveolar electrosensory cells has been the subject of active investigation, the actual mechanism by which the signals are first detected at the pores and traverse through the hydrogel within the canals still remains to be worked out [7, 8]. We and our collaborators have previously shown that the AoL hydrogel is extremely proton-conductive [9], and that this property may putatively contribute to the electrosensing mechanism by providing an electrical environment that is markedly more conductive than the surrounding seawater. The role that chitin might play in this overall scheme is unknown, in part because it is insoluble in aqueous solutions. However, by forming a composite material it may contribute to the gel’s space-filling properties (Fig. S1A,B) and serve as a scaffold for the interaction of proteins and sulfated glycosaminoglycans (proteoglycans) that are also components of the gel [9, 10]. Although we collected ample evidence of chitin in the AoL, preliminary X-ray diffraction experiments have suggested that the chitinous material may take on a different structure than chitin extracted from invertebrates. It is still unclear how chitin, an insoluble molecule, could exist as a hydrogel component, so the findings laid out here beg further investigation into the molecular structure of AoL gel.

## SUPPLEMENTAL INFORMATION

Supplemental Information includes two figures, supplemental experimental procedures and supplemental references.

## ACKNOWLEDGMENTS

We thank Michael Rego for help with data collection, Martin Cohn for catshark samples, Thomas Quinn, Jason Cope, Scott Hamilton, and Matthew Jew for ratfish specimens, Adam Summers and Tsutomu Miyake for embryonic cleared-and-stained skeletal preparations of embryonic stingray and skate, respectively, Andrew Gillis for technical advice and protocols for ISH, Larry Dishaw, John Cannon and Gail Mueller for the *Ciona* VCBP chitin probes, Eli Maciel for help with ISH and provision of planarian controls and probes, Marco Rolandi for advice on chitin structure, John Selberg for the keratan sulfate sample, Neil Hammerschlag for the tiger shark picture, Joel Sohn for procurement of a big skate specimen and early input on this work, and Murat Kaya for the cuttlebone chitin spectra and technical advice. We thank Biswa Choudhury at the UC San Diego GlycoAnalytics Core for help with the monosaccharide analyses. This work was funded, in part, by grants from the National Institutes of Health (R01-GM090049 to CTA), by discretionary funds from the Benaroya Research Institute and University of California-Merced, and by a University of California-Merced Senate award to CTA and LSH. Initial parts of this work were carried out at the Benaroya Research Institute, Seattle, Washington, USA.

