## Supplemental Information & Figures for "Evidence of Chitin in the Ampullae of Lorenzini of Chondrichthyan Fishes"

**SUPPLEMENTAL INFORMATION: Evidence of chitin in the Ampullae of Lorenzini of chondrichthyan fishes**

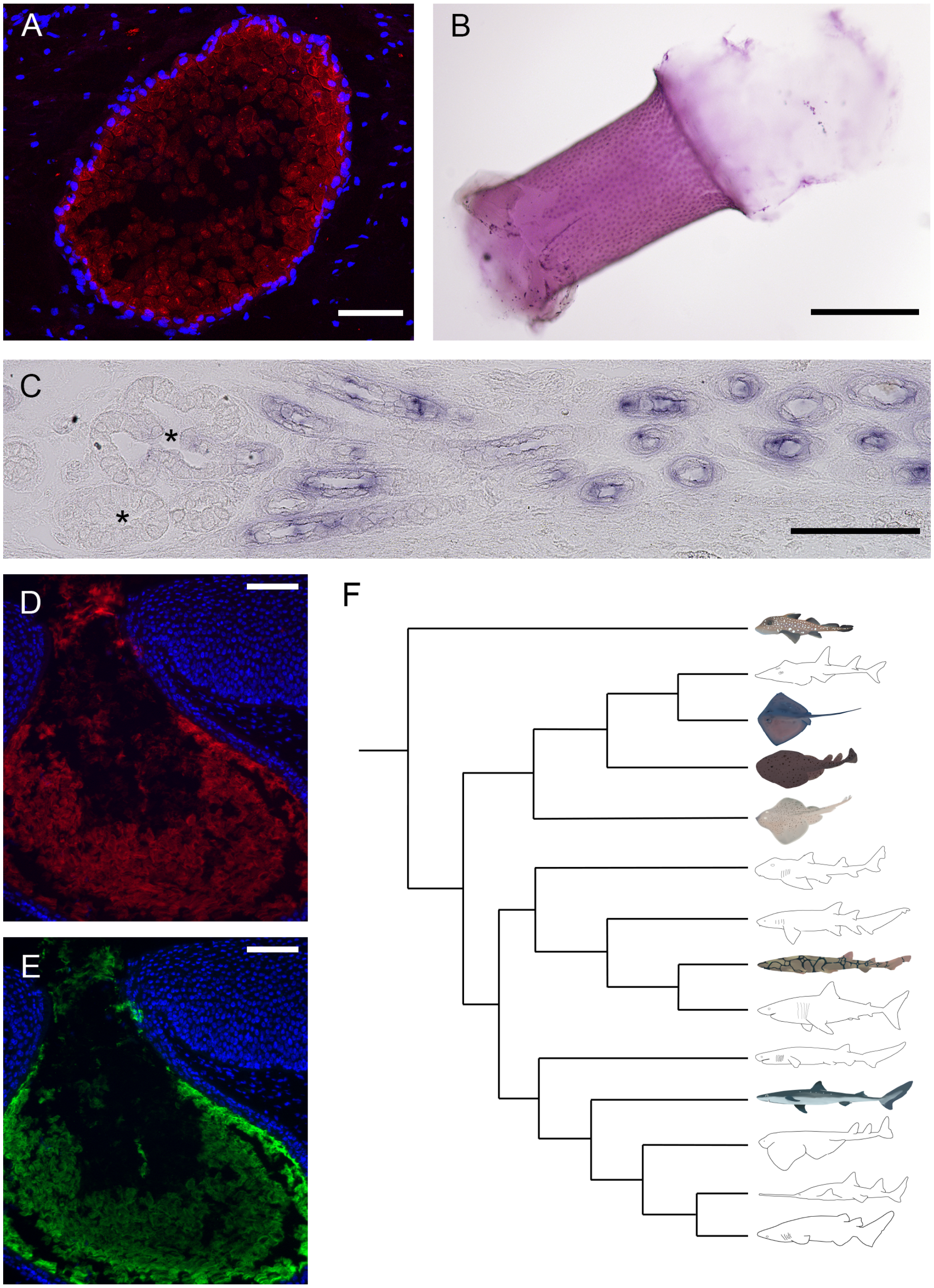

**Figure S1.** Chitin detection and characterization in the Ampullae of Lorenzini (AoL) of chondrichthyans. (A) Affinity histochemistry using tissue from a juvenile dogfish shark (*Squalus acanthias*) (0.4 m total length). Image of a 10 µm-thick cryosection through an AoL canal after affinity histochemistry (CBD-546, red; DAPI counterstain, blue). Note the absence of nuclei within the lumen of the AoL canal and the localization of CBD signals to the lumenal hydrogel. (B) Hematoxylin-Eosin staining of a short segment of an AoL canal from an adult big skate (*Raja binoculata*). This image shows the epithelial cells on the inside of the canal wall and the space-filling hydrogel spewing out of the tube. AoL hydrogel noticeably swells when such samples are placed in an aqueous solution (despite having been histologically fixed). (C) *In situ* hybridization with chitin synthase (*LeCHS*) riboprobe using a tissue section from an embryonic little skate (*Leucoraja erinacea*). *CHS* expression is observed within the cells that line the AoL canals, however, expression is absent from cells within the alveoli (as indicated by asterisks). (D, E) CBD labeling of a paraffin section from the snout of a spotted ratfish (*Hydrolagus colliei*) using fluorescent probes from two different chitin binding domains as described in [S1]. The CBD-546 probe used in D (red signal) was derived from a chitinase of the bacterium, *Bacillus circulans*, whereas the VCBP probe used in E (green signal) was derived from a chitin-binding immune receptor protein found in the gut of the tunicate, *Ciona intestinalis.* The probes were applied simultaneously to the same histological section and show highly similar fluorescent patterns. (F) A phylogeny depicting orders within the class Chondrichthyes represented by drawings of selected species. These are (from top to bottom): Chimaeriformes, Rhinopristiformes, Myliobatiformes, Torpediniformes, Rajiformes, Heterodontiformes, Orectolobiformes, Carchariniformes, Lamniformes, Hexanchiformes, Squaliformes, Squatiniformes, Pristiophoriformes, Echinorhiniformes. Clades portrayed by colored fish contain at least one representative species shown to possess chitin within their AoL using CBD histochemistry. The clades represented by black-and-white fish have not yet been examined. Because CBD signals were observed in the AoL of diverse chondrichthyan species, we hypothesize that chitinous gel emerged in a common ancestor to all chondrichthyan species. Phylogenetic positions of the orders shown in this phylogeny are based on gene sequence data from the 'Chondrichthyan Tree of Life" at sharksrays.org. Images were captured using the following microscopes: confocal microscope (A), stereomicroscope (B), compound light microscope (C), compound epifluorescence microscope (D, E). Scale bars are 200 µm (A), 300 µm (B), 100 µm (C), and 50 µm (D, E).

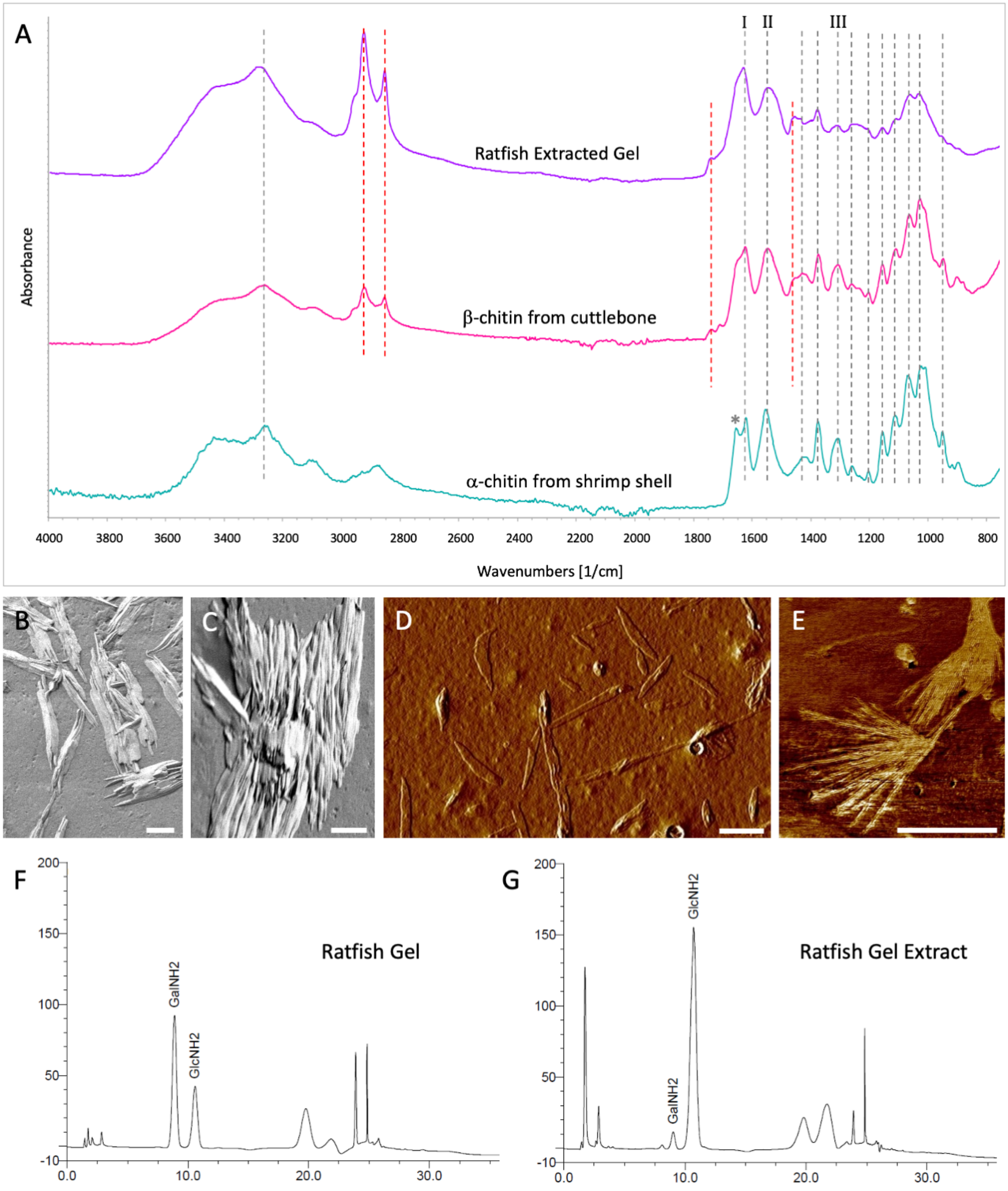


**Figure S2.** Physicochemical evidence of chitin in the AoL gel: analysis of polysaccharides extracted from the AoL gel of spotted ratfish, *Hydrolagus colliei*, by Fourier transform infrared spectroscopy (FTIR), scanning electron microscopy (SEM), atomic force microscopy (AFM), and monosaccharide analysis. (A) Comparison of FTIR spectra from AoL gel after treatment with

Proteinase K, HCl, NaOH, and extensive dialysis against water (purple), β-chitin obtained from cuttlebone of cuttlefish (pink), and α-chitin from shrimp shells (green). Black dotted lines indicate shared peaks among all three samples; red dotted lines indicate shared peaks among two of the samples. The peaks designated by I, II, and III are known as amide I, amide II and amide III peaks, respectively. The * at 1654 $cm^{-1}$ on the spectrum from shrimp shell chitin represents the split amide I band seen with the α- allomer [S2, S3]. We observed more similarities between the extracted ratfish gel and β-chitin from cuttlebone, but all three samples shared many of the same peaks. (B, C) SEM images of digested and extracted ratfish gel that had been dried down on a dish then scraped up and applied to a stub. We observed thick layers of material that were overlaid with crystals closely resembling chitin nanowhiskers [S4, S5]. (D, E) The same nanowhisker-like structures were observed with tapping mode AFM (D is an amplitude image, and E is phase). The two images show crystals of different shapes and scales obtained from two separate extraction experiments. The self-organization of crystal nanowhiskers into aggregates was readily observed (C, E), the degree of which seemingly attributable to the conditions at which the samples were prepared and dried before examination. (F, G) HPAEC-PAD profiles showing the monosaccharide content of ratfish gel and extracted ratfish gel after hydrolysis. The profile obtained from native ratfish AoL gel (F) showed obvious signals of glucosamine ($GlcNH_2$) and galactosamine ($GalNH_2$), as determined by $GlcNH_2$ and $GalNH_2$ standards that were run in tandem. It is unclear if $GalNH_2$ exists as a monosaccharide in native ratfish gel or if it is a component of a larger polymer. The profile resulting from analysis of the ratfish gel extraction (G) showed that measurable $GalNH_2$ was markedly reduced after the extraction treatment. However, the gel extract's profile did still show obvious $GlcNH_2$ signals, which suggests chitin was likely maintained in the sample after proteinase K digestion and after the extraction procedure. Scale bars are 1 μm.

### SUPPLEMENTAL EXPERIMENTAL PROCEDURES

#### Biological Specimens

Juvenile dogfish (*Squalus acanthias*) and adult skates (*Raja* spp.) were obtained from Carolina Biological Supply Company as formalin-fixed and preserved (Carosafe) specimens. Cleared-and-stained skeletal preparations of embryonic stingray (*Hypanus sabinus*) and little skate (*Leucoraja erinacea*) were obtained from Adam Summers (Friday Harbor Laboratory) and Tsutomu Miyake (Benaroya Research Institute), respectively. Martin Cohn (University of Florida) provided several fixed specimens of embryonic and juvenile specimens of the spotted catshark (*Scyliorhinus canicula*). A spotted ratfish (*Hydrolagus colliei*) was collected by Thomas Quinn (University of Washington) on a research cruise in Puget Sound, and brought to the Benaroya Research Institute for processing. Additional expired and frozen spotted ratfish were obtained from Jason Cope (NOAA), Scott Hamilton (Moss Landing Marine Labs, MLML) and Matthew Jew (MLML). Pacific electric ray (*Tetronarce californica*) specimens were collected off the coast of Monterey Bay by John Field (NOAA Southwest Fisheries Center) and his NOAA groundfish assessment team. Specimens of live embryonic and adult little skates were obtained from the Marine Resources Center (Marine Biological Laboratory). All animal work was conducted under approved IACUC protocols: IACUC16-014 (Benaroya Research Institute) and AUP18-0001 (University of California, Merced). Euthanasia was carried out using tricaine methylsulfonate (Western Chemical Inc., #NC0872873) at 1000 mg/L.

**Paraffin Sections**

Wax sections of formalin-fixed embryonic or adult tissues were prepared using standard processing, embedding and microtomy protocols. If the sections were to be used for *in situ* hybridization, RNase-free reagents were used in the processing.

**Cryosections**

Fixed tissues were washed in PBS (3 x 10 minutes) and equilibrated in 15% sucrose in PBS for three hours at room temperature. The solution was replaced with 15% sucrose/7.5% gelatin in PBS, and samples were incubated for three hours at 37 C. Samples were then infiltrated with 20% gelatin in PBS overnight at 37 C and embedded in fresh 20% gelatin in PBS using plastic molds. Embedded samples were mounted onto a cryotome chuck with Tissue-Tek O.C.T. compound, flash frozen in liquid nitrogen and sectioned at 10-12 µm on a cryotome and mounted on Superfrost Plus slides (VWR). Sections were de-gelatinized with 0.3% gelatin in 50% ethanol and allowed to completely air dry before histochemistry and staining.

**Affinity histochemistry probes**

The two chitin probes used in this study were selected on the basis of availability and known efficacy of chitin binding. Most of the experiments used a probe that was obtained from New England Biolabs because of the ease of handling (one step method):

AWQVNTAYTAGQLVTYNGKTYKCLQPHTSLAGWEPSNVPALWQLQ (45 aa)

Chitin binding domain from plasmid pYZ205 (*Bacillus circulans*)
Conserved domain superfamily cluster cl00046 (NCBI)

We also used a different chitin probe in order to independently validate the presence of chitin:

FTCEGKADGLYSDPYQCNMYYECVMYVKYHRPCPTGTVF (39 aa)
Chitin binding domain from VCBP_C (*Ciona intestinalis*, NP_001190979)
Conserved domain superfamily cluster cl02629 (NCBI)

**Isolation and labeling of SNAP-tag Chitin Binding Domain (SNAP-CBD) fusion protein**

Plasmid pYZ205 containing the SNAP-CBD fusion construct was provided in T7 Express *E. coli* cells and protein isolated on a nickel-NTA agarose resin affinity column as described in Maduzia et al. [S6]. The protein was eluted in 1 mL fractions with elution buffer (50 mM $NaH_2PO_4$, 300 mM NaCl, 250 mM imidazole, pH 8.0), quantified on a Nanodrop spectrophotometer, and electrophoresed by SDS-PAGE to ascertain purity. Fractions with the highest concentration of protein were pooled and dialyzed overnight in phosphate buffer (50 mM $NaH_2PO_4$, 50 mM $Na_2HPO_4$, pH 7.2, 0.1 mM NaCl) containing 1 mM DTT, then stored at -80 C. To fluorescently label the fusion protein, 5 μM SNAP-CBD was incubated with 10 μM SNAP-Surface Alexa Fluor-488 or -546 (New England Biolabs) and 1 mM DTT in phosphate buffer for one hour at 37 C in the dark, then dialyzed overnight at 4 C in phosphate buffer with 1 mM DTT. An aliquot of the fluorescently labeled SNAP-CBD was electrophoresed by SDS-PAGE and imaged with a Typhoon 9410 Imager (GE Amersham Molecular Dynamics) to ensure successful labeling, then stored at -20 C in 50% glycerol. In order to detect chitin *in situ*, we employed affinity histochemistry using this CBD peptide probe (CBD-488 or CBD-546).

Alternatively, aliquots of the SNAP-CBD were labeled with biotin using either SNAP-surface biotin (as above) or through direct conjugation. For the latter, approximately 0.2 mg of SNAP-CBD was labeled with NSH-$PEG_4$-Biotin with EZ-Link Micro NSH-PEG4-Biotinylation Kit

(Thermo Scientific), which attaches NSH-PEG$_4$-Biotin to lysine residues and the N-terminus. Unincorporated NSH-PEG$_4$-Biotin was removed by desalting columns in the Kit. Biotin incorporation was measured using Biotin Quantitation Kit (Thermo Scientific); only samples showing incorporation of one or more biotin per molecule of the SNAP-CBD were used for further experiments.

**Chitin affinity histochemistry**

Chitin affinity histochemistry largely followed the protocols outlined in Tang et al. [S1]. Fixed specimens (whole-mount tissue or sectioned material) were washed in PBS-T (PBS with 0.05% Tween-20) at room temperature (3 x 10 minutes), then dehydrated in a graded methanol series and stored at -20 C. For histochemistry, specimens were rehydrated with PBS-T, bleached briefly with 1% $H_2O_2$/0.5% KOH at room temperature to remove pigment, then permeabilized with 1% Triton-X/PBS overnight at 4 C. Specimens were then washed in PBS-T for thirty minutes, and blocked in 1% BSA in PBS for one hour before incubation with fluorescently-labeled CBD-488 or CBD-546 at 4 C for 24-48 h. For co-detection experiments, specimens were simultaneously incubated with either *Ciona intestinalis* VCBP-C [S7] fused with the C-terminal region of human IgG1 Fc (provided by Larry Dishaw, John Cannon and Gail Mueller, University of South Florida) or CBD-biotin. For secondary detection of VCBP-C or CBD-biotin, larvae were washed in PBS-T (3 x 10 minutes), blocked for one hour in 1% BSA in PBS or Pierce TBS Blocking Buffer (Thermo Fisher, 37535), then incubated overnight at 4 C with goat anti-human IgG Fc conjugated to Dylight 488 (Abcam) or Streptavidin-488 (Invitrogen), respectively. Samples were then washed in PBS-T (3 x 1 hour), counterstained and imaged. If using slides, samples were mounted in anti-fade mounting medium (Vectashield) prior to imaging.

**Chitinase digestion of AoL-bearing histological specimens**

Slides containing histological sections were rehydrated in 1X PBS. Several types of chitinase were used but we found that chitinase isolated from the bacterium, *Streptomyces griseus* (Sigma Aldrich, C6137) worked most reliably. Chitinase was diluted to 1 mg/mL in 50 mM $KH_2PO_4$ and 10% glycerol. Slides were incubated in 300 μL of chitinase solution under a piece of parafilm and placed in a humidified chamber at room temperature for approximately 36 h. Samples were washed 3 times for 30 minutes in 1X PBS and then 300 μL of Pierce TBS Blocking Buffer were added to the slides and incubated for 1 h at room temperature. Slides were stained with CBD-546 diluted 1:40 in TBS Blocking Buffer at 4 C overnight. Slides were washed 3 times in 1X PBS and then counterstained with a dilute DAPI solution (in 1X PBS). Slides were washed 2 times in 1X PBS and then rinsed with $dH_2O$ and mounted with anti-fade mounting medium (Vectashield) before imaging.

**Pilot transcriptome of the AoL of adult little skate**

Approximately 600 mg of alveoli and canal tissue from the hyoid cluster was dissected from an adult little skate and stored in RNAlater (Life Technologies, #AM-7020) at 4 C. The tissue was homogenized, and total RNA was isolated using Trizol reagent (Life Technologies, #15596-026) and purified with RNeasy MinElute Cleanup kit (Qiagen, #74204) according to manufacturer's instructions. The library was prepared and sequenced by Otogenetics Corporation (Atlanta, GA) using an Illumina HiSeq 2500 sequencer in Rapid Run mode with SBS v2 chemistry. The raw paired read files were quality-trimmed with Trimmomatic and assembled into a *de novo* transcriptome using Trinity v. 2.2.0. The sequence data were submitted to the SRA database under the BioProject PRJNA550453 (SRA Experiment SRX6358492).

**Probe synthesis and *in situ* hybridizations**

RNA was extracted from *L. erinacea* tissue and cDNA was synthesized from the extracted RNA using the SuperScript III First-Strand Synthesis kit from Invitrogen (Life Technologies, #18080-051). Gene-specific primers were designed for a 645 bp segment of a little skate chitin synthase gene identified using both our little skate transcriptome (above) and from [S8]. This probe, *LeCHS*, is given below (magenta sequences are the PCR primer sequences).

```
CCACCAAAAGGAGTTCTAGCCATGGTATGCTTCATCGAAGCAATTATCGCACTTGCCACTTCAGTTCTGGTGCTGGTCTGCTTGCC
TCAGTTTGATGTCATCACAAATCTCTGCATTTTGAATGGTGCCTGTCTGTTTCCTTCATTCCTACAAATAATGTATGAGTTGAAAC
ATCTGGGGGTTGCCTTGCTATTCCCCCTCATTGGTTTTGGGTTGACTTTTCTAGGCCTCTTGCTTTTCATTTTAGTCCACAATTCC
CTGCAGCAATCAGACACCCCTCAATACCTTCAGATGTACGTAGCCATAGCAATAGTTTCCTTAACAGTATTATCTCTCAATTGGTG
GGAAAACTTTACATCCTTCTGTAAAATTGGATTCCTGCAAAATATAAGGGAAAGCCTACGTGGCAATAGAAATCTTACCTACGTTT
GCAGTAGTGCAGTTAGAATTCTCGTCACCTTTGGTGTAGTCGCGGCATGGATTCCAATTAAGAAGTATGACTGGGTAGATTTGAAA
ACAGTCTCTCAATTTGAACTAAATATTGTTTTGGGTCTCTTTGGGGTGCAGGCATGTTCTTCTGTACTTTGCCACTGGTTTGGGGT
CCTGGTGTGCAAAATGCATGCAGTCAGGCGAAGTTTCATGCTG
```

The fragment was cloned into a pCR-II-TOPO vector (Invitrogen, #450640) and DIG-labeled probes were synthesized using Ambion's MEGAscript Sp6 and T7 kits (Invitrogen, #1330 &

#1334). Probes were stored in hybridization buffer (50% Formamide, 5X SSC, 2% blocking reagent (BBR), 50 μg/mL heparin, 10 μg/mL yeast tRNA, 0.1% CHAPS, 0.1% Triton-X) at a concentration of 100 ng/μL at -80 C. The planarian ChAT probe was obtained from Eli Maciel at UC Merced for use as a negative control (see below).

Embryos were fixed in HA fixative (70% ethanol, 5% glacial acetic acid, 4% formaldehyde, 21% water) overnight then moved to RNAse-free methanol and stored at -20 C until use. Snout and fin pieces were dissected from embryos for whole-mount ISH experiments and entire anterior ends were embedded in paraffin for ISH with sections.

For whole-mount ISH, we adapted and modified the protocol from Andrew Gillis (Cambridge University). Embryos were moved to 1X PBT. To permeabilize, the tissues were incubated in proteinase K (5 μg/mL) for 15 minutes then in glycine (2 mg/mL) for 5 minutes. The tissue was re-fixed in fresh 4% paraformaldehyde (PFA) for 20 minutes. Hybridization buffer (hyb) was added to each tube for ten minutes then replaced with fresh pre-warmed hyb (68 C) and incubated for 3-5 hours. Probes were diluted to a working concentration of 5 ng/μL in 1 mL hybridization buffer and then denatured for 8 minutes at 80 C. Pre-hybridization buffer was removed, replaced with probe solution, and the tubes were incubated at 68 C for 70 hours.

2X SSC (10 mL formamide, 4 mL 20X SSC (pH 4.5), 2 mL 10% SDS, 4 mL $dH_2O$), and 0.2X SSC (10 mL formamide, 2 mL 20X SSC (pH4.5), 8 mL $dH_2O$) were pre-warmed to 68 C. Probe was removed, 2X SSC was added quickly to the tissue and washed for 45 minutes at 68 C (this wash was performed two times). Three 0.2X SSC washes were then performed at 68 C for 30 minutes each. Tissue was then washed 4 times in MABT (Maleic Acid Buffer + Tween-20) for 10 minutes each. Fresh blocking solution (20% heat inactivated sheep serum + 1% Roche blocking reagent) was prepared and added to the tissue for 4 hours at 4 C. Blocking solution was then replaced with anti-DIG AP Fab fragments (Roche, #11093274910) diluted 1:2500 in blocking buffer. Tubes were left at 4 C overnight with gentle rocking.

The next day, samples were washed 3 times in MABT for 5 minutes each and then 5 more times in MABT for one hour each. The samples were left in the last MABT wash overnight at 4 C. Samples were then washed in freshly made NTMT (1 mL 5 M NaCl, 5 mL 1M Tris/HCl, 0.2 mL Tween-20, 44 mL $dH_2O$) three times for 10 minutes. Tissue was bathed with BM Purple solution (Roche, 11442074001) rotating in the dark at room temperature. Staining solution was replaced after an hour. After the color reaction was complete, tissues were washed with 1X PBS then post-fixed with 4% PFA and stored in 75% glycerol.

For ISH with sections, slides were rehydrated in 1X PBT then rinsed with 2X SSC in a Coplin jar at room temperature. DIG-labeled probes were diluted to a working concentration of 5 ng/μL in slide-specific hybridization buffer (50% formamide, 1X salt solution, 10% dextran sulfate, 10 mg yeast tRNA, 1X Denhardt's solution, and DEPC water to 10 mL). The working concentration of probe (5 ng/μl) was denatured at 80 C for 5 minutes before adding to each slide and covering with a #1 coverslip. Slides were placed in a humidified chamber of 50% formamide, 50% 2X SSC and incubated overnight at 63 C.

The next day, slides were washed twice with a solution of 50% formamide, 1X SSC, and 0.1% Tween-20 at 63 C for 30 minutes. Slides were washed 3 times with 1X MABT solution, placed in blocking solution (1% Roche blocking reagent and 20% sheep serum) for 2 hours, then incubated in anti-DIG-AP antibody at a concentration of 1:1000 in block solution overnight.

Slides were washed every 30-60 minutes with MABT the following day and left in the final wash overnight at 4 C. On the final day, fresh NTMT (0.1 M NaCl, 0.1 M Tris (pH 9.5), 5 mM $MgCl_2$, 0.1% Tween-20) was prepared and used to wash the slides 3 times. Slides were incubated in BM Purple on a shaker for several hours until color was revealed. In some cases, staining continued overnight in order for signal to be observable. Slides were washed with 1X PBS, post-fixed for one hour with 4% PFA then mounted with glycerol or antifade medium (Vectashield) for imaging.

Sense probes were synthesized in tandem with antisense *CHS* probes as described above. Upon performing ISH experiments with sense probes, we often observed localization of signal to the AoL in the same specific pattern that was seen with the antisense probes. Other researchers have reported similar observations when performing *in situ* hybridization on chondrichthyan species, presumably due to an abundance of antisense transcripts. We thus used heterospecific probes from another organism as our negative control. We used an antisense probe specific to a cholinergic acetyltransferase (*ChAT*) gene of planaria (*Schmidtea mediterranea*, SMU15019525 in SmedGD database) (complements of Eli Maciel, Oviedo lab, UC Merced). When we performed ISH experiments with *LeCHS* probe using skate tissue, we would use either a second piece of tissue from the same skate specimen or another slide in the case of tissue sections, but substitute *LeCHS* for *ChAT*. When we did whole-mount ISH experiments, we would also use whole planarians in tandem with skate tissues to confirm that *ChAT* probes hybridized to the expected anatomic locations in the flatworms.

**Imaging**

Stereoscope imaging was done on a Leica M205FA fluorescent stereoscope equipped with a DFC360FX monochrome CCD camera and a DFC425C color CCD camera. Epifluorescent images were taken using either a Leica DMR upright epifluorescent microscope equipped with a SPOT RT Slider cooled 1.4 megapixel color/monochrome CCD camera and an Insight 4 megapixel color CCD camera (Diagnostic Instruments) or a Keyence BZ-X700 microscope workstation equipped with epifluorescence. The latter was also employed for bright field images. Confocal images were obtained with a Leica TCS SP5 laser scanning confocal microscope. Brightness and contrast adjustments, gamma correction, background subtraction, and confocal image smoothing using the unsharp mask filter were completed in Photoshop CS9 (Adobe) and Fiji (ImageJ, NIH).

**Collection of gel from the Ampullae of Lorenzini of *Hydrolagus colliei***

Isolation of AoL gel was previously described [S9]. Briefly, gel was obtained from multiple freshly caught and recently expired fish by gently pressing on the visible surface pores and collecting the extruded material using a mechanical plunger-style pipette or by scooping it directly into tubes with a pipet tip or scalpel. Samples were stored at −80 C until use.

**Polysaccharide extraction and analysis by Fourier transform infrared spectroscopy (FTIR), scanning electron microscopy (SEM), and atomic force microscopy (AFM)**

Several milliliters of gel were squeezed from the AoL pores of expired spotted ratfish (*Hydrolagus colliei*). We carried out a few separate extraction experiments and changed the protocol slightly each time. In one of the extractions, gel was diluted in an equal volume of Proteinase K buffer (0.1M Tris, 0.05M EDTA, 0.1% SDS). Proteinase K (Roche Cat No.3115879001) was added to a final concentration of approximately 40 µg/mL and the solution was incubated at 50 C for 1-2 h. We then brought the sample to 75 C for 20 minutes to denature remaining Proteinase K. The sample was moved to a Pur-A-Lyzer Mega 3500 dialysis unit (Sigma Cat No.PURG35015-1KT) and dialyzed against water overnight. In our other extraction rounds, we did not use a Proteinase K treatment, but added the gel directly in the dialysis unit and dialyzed against water overnight.

Samples were heated to 50 C in water with gentle spinning then added to pre-heated 1M HCl and allowed to incubate for 1 h at 50 C. Sample was brought slowly to room temperature then dialyzed against several rounds of water over the course of a day and overnight. Sample was brought again to 50 C in water, then incubated in 4-7 rounds of 1 N NaOH at 50 C for 1-2 h each time. Sample was dialyzed in many changes of water for several days until pH reached approximately 6.5-7. For the extraction round that did involve a Proteinase K step, a subset of the resulting liquid was placed in an Amicon Ultra 100 kD centrifugal filter unit (Millipore Sigma Cat No.UFC510024) and spun-dialyzed with 6 washes of water to remove any small proteins, sugars, or other molecules smaller than 100 kD. Samples were dropped onto glass slides and dried overnight at 62 C before being used for FTIR. For scanning electron microscopy (SEM) and atomic force microscopy (AFM), samples were either dropped directly onto mica (AFM), or dropped on a petri dish, scraped up and affixed to an aluminum stub using carbon tape (SEM).

Chitin from *Sepia sp*. cuttlebone (from pet store) was isolated using an established protocol [S3]. Cuttlebone (β-) chitin, shrimp shell (α-) chitin (Sigma, C9752) and polysaccharide extract from ratfish gel (three separate isolations) were dried at 62 C overnight before spectra were recorded with a Bruker Vertex 70 FTIR spectrometer with a diamond crystal ATR accessory. In addition, requisite quality control FTIR experiments were also carried out with keratan sulfate [S10] and chitosan (75% deacetylated chitin; Sigma, C3646) in order to validate the robustness and fidelity of our AoL chitin extraction method. Raw data for all FTIR spectra in this project are available at: https://drive.google.com/open?id=1PRKNVqg2nmxJq3M8y-MSjzsBf6KfmFYe.

Scanning electron microscopy was performed with the Zeiss Gemini500 FEG-SEM at the Imaging and Microscopy Facility of University of California, Merced. Ambiently dried particles were adhered to conductive carbon tape. Beam landing energy of 500 eV and secondary electron detection were used.

Atomic force microscopy (AFM) was performed using a Veeco Innova instrument microscope in tapping mode. Probes with a spring constant of 40 N/m (Tap300Al-G, Budget Sensors) were used to image air-dried samples drop-cast on freshly cleaved mica to obtain topographical, amplitude and phase images of crystals formed on the surface in air. Images were collected at a scan rate of 1 line per second at room temperature.

**Monosaccharide analysis with ratfish gel and extracted polysaccharides from gel**

Monosaccharide analysis of ratfish native gel and extracted gel was performed by the UC San Diego GlycoAnalytics Core. Samples of shrimp chitin (Sigma, C9752), spotted ratfish (*Hydrolagus colliei*) gel, and extracted ratfish gel (see section 1.3.13 for extraction protocol) were sent to Biswa Choudhury at the GlycoAnalytics Core. Approximately 0.2 mg of shrimp chitin control was weighed out in a glass hydrolyzing tube and hydrolyzed. 1.5 mg lyophilized ratfish gel was dissolved in 750 µL of ultrapure water and homogenized to fine suspension. 100 µg of material was taken for hydrolysis. The stickiness of the extracted ratfish gel prevented it from being weighed so it was dissolved in 500 µL ultrapure water (clear solution) and 50 µL was taken for hydrolysis (10% of the total material). All samples were hydrolyzed with 6 N HCl at 100 C for 6 hours, followed by removal of acid by dry nitrogen flush. Finally, dried and hydrolyzed samples were dissolved in water and approximately 0.25 µg of chitin control, 3 µg of ratfish gel, and 0.625% of ratfish gel extract were injected on HPAEC-PAD. Known amounts of standards were used for quantification of the monosaccharides.

AUTHOR CONTRIBUTIONS

M.P., W.J.T. and C.T.A. conceived the original experiments; M.P., W.J.T., M.R., D.O.D., K.H., and C.T.A. collected the data and conducted the analyses with input from all authors. M.P. and C.T.A. wrote the paper with input from all authors.

DECLARATION OF INTERESTS

The authors declare no competing interests.
